# Susceptibility of Cu(I) transport ATPases to sodium dodecyl sulfate. Relevance of the composition of the micellar phase

**DOI:** 10.1101/2023.03.13.532330

**Authors:** Alvaro A. Recoulat Angelini, Noelia A. Melian, F. Luis González-Flecha

**Affiliations:** Universidad de Buenos Aires - CONICET, Laboratorio de Biofisica Molecular. Instituto de Química y Fisicoquímica Biológicas, Argentina

**Keywords:** P-ATPases, Inactivation, Detergents, SDS

## Abstract

Sodium dodecylsulfate (SDS) is a well-known protein denaturing agent. A less known property of this detergent is that it can activate or inactivate some enzymes at sub-denaturing concentrations. In this work we explore the effect of SDS at sub-denaturing concentrations on the ATPase activity of a hyper-thermophilic and a mesophilic Cu(I) ATPase reconstituted in mixed micelles of phospholipids and a non-denaturing detergent. We first develop an iterative procedure to evaluate the partition of SDS between the aqueous and the micellar phases. This procedure allows to determine the composition of micelles prepared with variable amphiphiles content. When incubating the enzymes with SDS in the presence of different amounts of phospholipids, it can be observed that higher SDS concentrations are required to obtain the same degree of inactivation when the initial concentration of phospholipids is increased. Notably, we found that, if represented as a function of the mole fraction of SDS in the micelle, the degree of inactivation obtained at different amounts of amphiphiles converges to a single inactivation curve. To interpret this result, we propose a simple model involving active and inactive enzyme molecules in equilibrium. This model allowed us to determine the Gibbs free energy change for the inactivation process and its derivative respect to the mole fraction of SDS in the micellar phase, this last being a measure of the susceptibility of the enzyme to SDS. Our results showed that the inactivation free energy changes are similar for both proteins, and indicate that the equilibrium is highly shifted towards the active form in both enzymes. Conversely, susceptibility to SDS is significantly lower for the hyperthermophilic ATPase, suggesting an inverse relation between thermophilicity and susceptibility to SDS.

## 1. Introduction

Cu(I) transport ATPases constitute a group of integral membrane proteins involved in active transport of Cu(I) across cell membranes coupled to the hydrolysis of ATP [1]. They belong to the P-type ATPases family, all sharing some common structural features [2] including a transmembrane region -composed by 6-8 α-helices arranged in a helix bundle- and three cytoplasmic domains: the catalytic one, composed by subdomains P (phosphorylation) and N (nucleotide binding), the so-called actuator (A) and a metal binding domain at the N terminal extreme (N-MBD) (Fig. 1). As in all integral membrane proteins, highly hydrophobic residues in the transmembrane region face to the external surface interacting with phospholipids [3]. These interactions are critical for the consolidation of the three-dimensional structure and the enzymatic hydrolysis of ATP taking place in the catalytic domain [4].

**Fig. 1.**
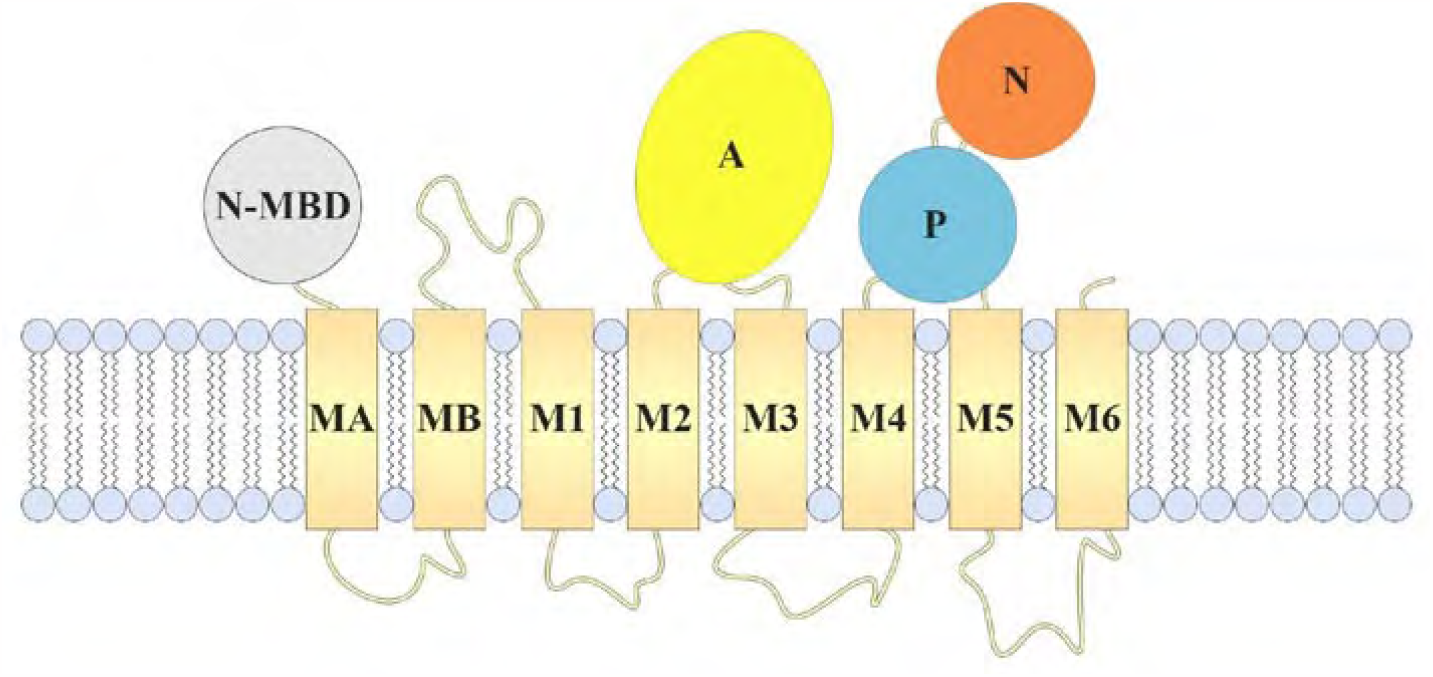
Schematic representation of Cu(I) transport ATPase structure. The transmembrane domain is composed by 8 α-helical segments, 6 are common to all P-type ATPases (M1 to M6) and 2 are specific to the P1B subfamily (MA and MB). The cytoplasmic domains are the catalytic one (composed by the nucleotide binding (N) and the phosphorylation (P) units), the actuator (A) and a metal binding domain (MBD) at the N-terminal extreme (some members such as *Af*CopA posses an additional MBD at the C-terminal extreme).

Sodium dodecysulfate (SDS) is a strong anionic detergent with known denaturing capacity at millimolar concentrations [5-7]. The underlying denaturation mechanism shows similar features to classical chaotropes like urea or GdnHCl and is driven by its higher global binding affinity for the denatured state than for the native state [6, 7]. However, the binding mechanism is different where the entropic contribution is the main driving force, showing increasing efficiency when compared to chaotropes [8]. Furthermore, sub-denaturing concentrations of this detergent have been reported to activate (e.g. [9]) or inactivate (e.g. [10]) some enzymes.

SDS was also proposed to quantify the kinetic stability of hyperstable proteins via a procedure named “time dependent SDS trapping” [11]. This procedure was based on incubating a protein with high SDS concentrations at high temperature as a function of time. The authors found that kinetics of the loss of SDS resistance correlated linearly with their unfolding rate. To date, there are no studies that determine whether a similar relationship holds at low concentrations of SDS.

In the presence of lipid membranes or micellar systems, SDS will partitionate between the aqueous and the lipid phases. Therefore, for membrane proteins reconstituted in mixed micelles or bilayers, this partition will determine how much SDS is available for interaction at the transmembrane surface (exchanging with lipids and non-denaturing detergents) and/or at the water soluble domains (interacting through the monomeric form of SDS present in the aqueous phase).

In this work we explored the effect of SDS at sub-denaturing concentrations on the ATPase activity of two Cu(I) ATPases belonging to the hyperthermophilic archaea *Archaeoglobus fulgidus* (*Af*CopA) [12] and the mesophilic bacteria *Legionella pneumophila* (*Lp*CopA) [13].

## 2. Materials and methods

### 2.1. Protein expression and purification

CopA cDNAs were cloned and transformed in *Escherichia coli* as described previously [13, 14]. Membranes isolated from *E. coli* cells expressing CopA were solubilized with C_12_E_10_. Protein purification was performed by affinity chromatography using a Ni^2+^-nitrilotriacetic acid column. Protein concentration was determined using the molar extinction coefficient for the purified proteins calculated from their sequence [15] after correcting by scattering the absorption spectrum of the purified enzyme. Proteins were stored frozen until use.

### 2.2. ATPase activity assays

ATPase activity was measured at the working temperature of each enzyme (70°C for *Af*CopA, and 37°C for *Lp*CopA) as the initial rate of release of inorganic phosphate (Pi) from ATP, in a medium containing 0.8 μM purified enzyme, 25 mM Tris-HCl (pH 7.0), 3 mM MgCl_2_, 2.5 mM sodium azide, 2.5 mM ATP, 20 mM Cys, 10 mM NaCl, 200 μM CuSO_4_, 2.5 mM DTT, and the asolectin, C_12_E_10_ and SDS concentrations indicated in each figure. Released Pi was determined using a malachite green procedure with some modifications [16, 17].

### 2.3. Critical micelle concentration measurements

Calorimetric demicellization experiments were performed as described by Tso et al [18] using a VP-ITC microcalorimeter (Micro-Cal, USA) at temperatures between 290 and 340 K. The sample cell was filled with 10 mM NaCl, 25mM Tris pH 7.0 at each working temperature, and detergent at high concentration in the same buffer were added into the sample cell step by step through the syringe.

### 2.4. Data analysis

At least three independent experiments were performed for each condition. The equations were fitted to the experimental data using a non-linear regression procedure based on the Gauss-Newton algorithm [19]. The dependent variable was assumed to be homoscedastic (constant variance), and the independent variable was considered to have negligible error. To select the best model to fit the experimental data we used the Second-Order Akaike Information Criterion (AICc). The best model was considered to be the one which gave the lower value of AICc [20]. Joint confidence regions for the regression parameters were calculated as described previously [21, 22].

## 3. Results and discussion

### 3.1. Determination of the critical micelle concentration of C_12_E_10_ and SDS

A calorimetric procedure was used to determine the cmc of each detergent in buffer (cmc°). For this task, a concentrated solution of each detergent (well above the expected value for the cmc°) in 10 mM NaCl 25 mM Tris HCl pH 7 at the working temperature was injected into the same buffer contained in the calorimeter cell. During the first injections two processes simultaneously occur, dilution of micelles in buffer, and demicellization, thus resulting in a diluted solution of monomeric detergent in buffer. When the concentration of detergent in the cell equals the cmc° value, only the first process will occur, i.e. the dilution of micelles in a solution containing the buffer and monomeric detergent at a concentration that will remain constant throughout the subsequent injections. This behavior is visualized, when the detergent was C_12_E_10_, as two different trends in the thermogram, each one characterized by a near linear dependence of the normalized heat as a function of the total concentration of detergent expressed as monomer. The intersection between both straight lines is considered to be the cmc° (Fig 2. A)

**Fig. 2.**
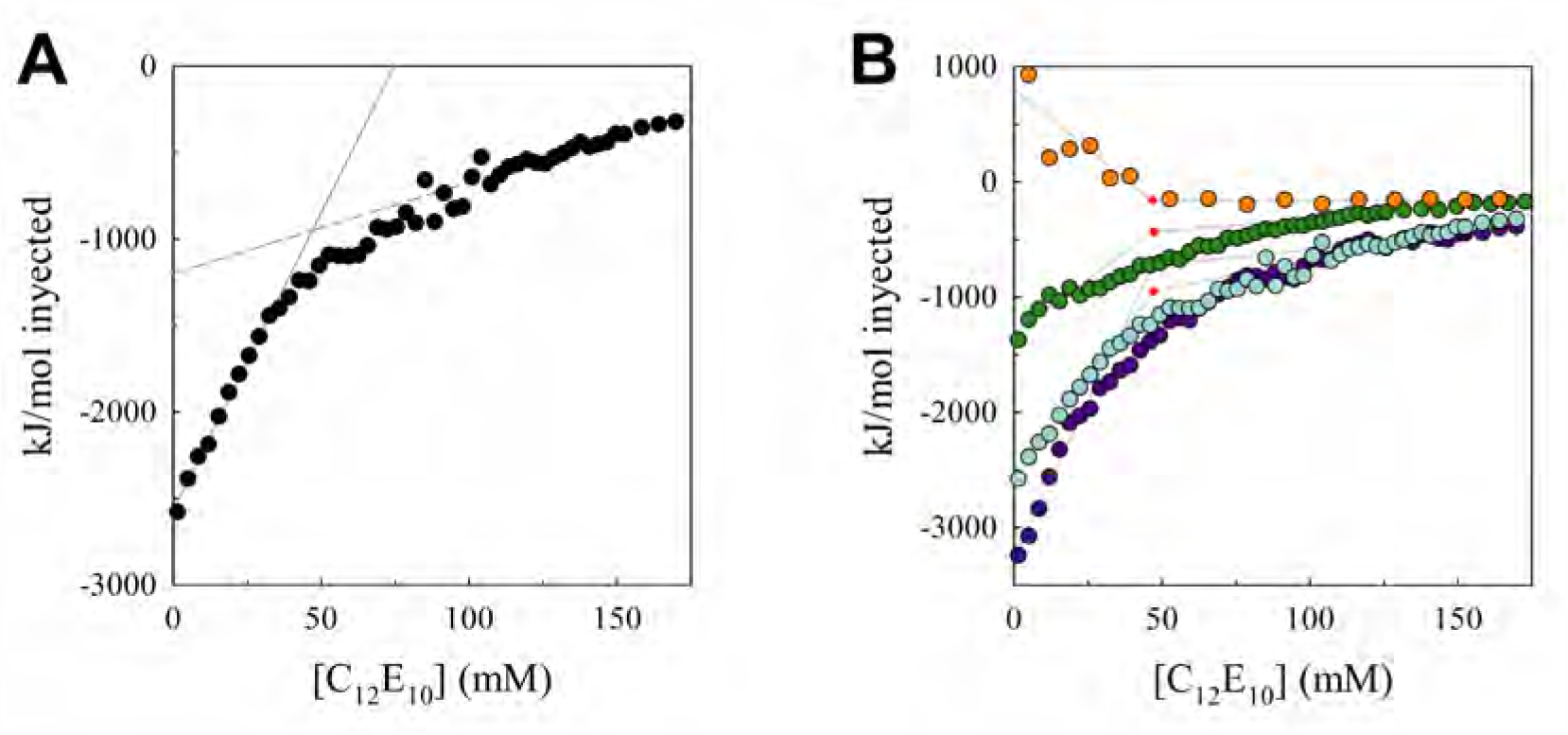
Thermogram for the titration of buffer with C_12_E_10_. (A) The heat per mole of injected detergent at 25 °C is represented as a function of the final concentration of detergent in the cell. For calculation of the cmc° two lines were adjusted to the regions of the graph where the change follows a linear function. The intercept represents the cmc° value. (B) The same procedure was repeated at the following temperatures: 15°C (purple), 25°C (light blue), 37°C (green) and 60°C (orange).

It can be observed that the intersection occurs at similar [C_12_E_10_] values for all the assayed temperatures (Fig. 2B).

In the case of SDS, the observed behavior is quite different. Below and above the cmc° there is a linear dependence of the exchanged heat per mole of detergent with a very small dependence on SDS concentration. Between the two linear regimes, a sigmoid region becomes evident in the thermogram. This behavior suggests that it is not a simple phase transition involving two well defined states, but a gradual transition between the monomeric region and the micelles region. It is possible that sigmoidicity is indicative of the formation of premicellar aggregates coexisting with monomers and full size micelles [23]. Given this particularity, the cmc° value is estimated as the inflection point in the sigmoid region (Fig. 3A)

**Fig. 3.**
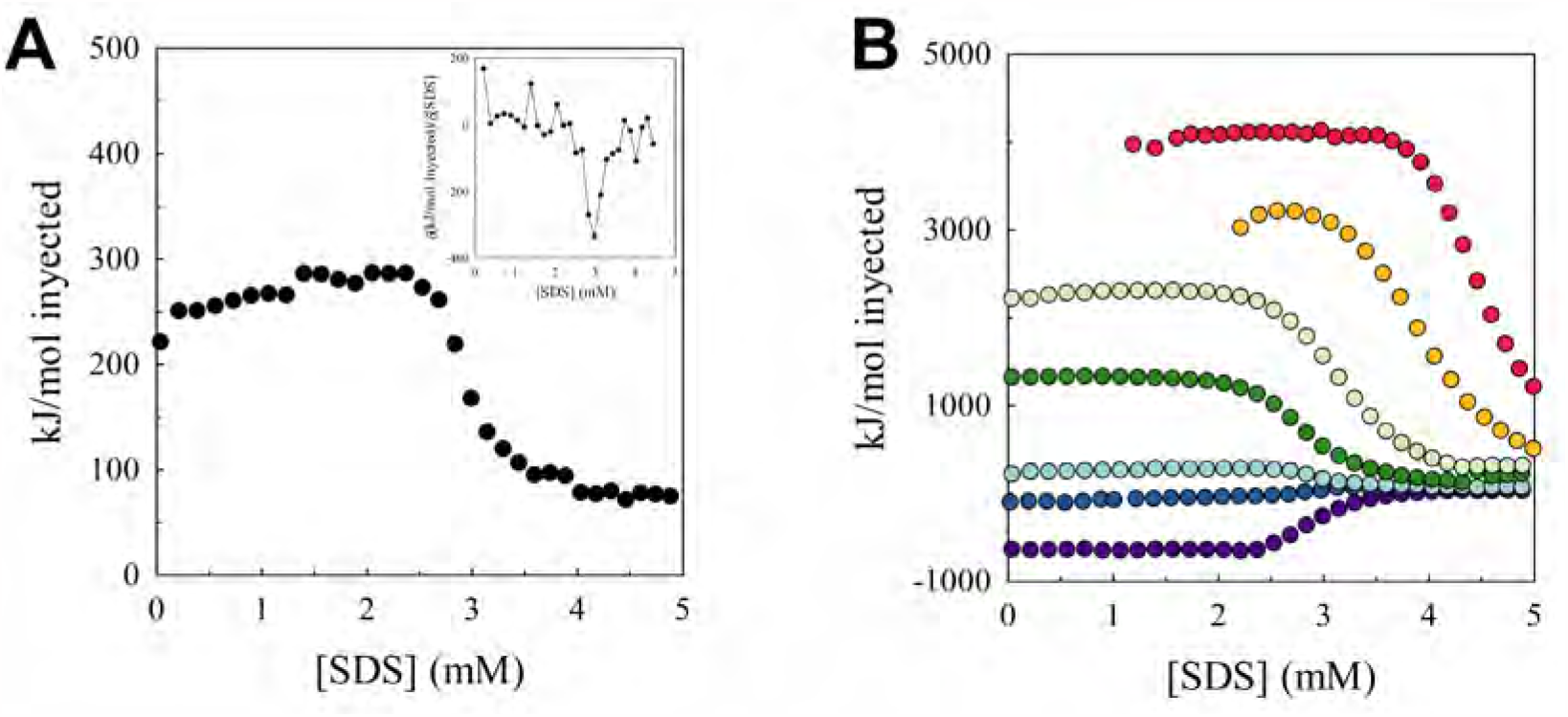
Thermograms of the titration of buffer with SDS. (A) The exchanged heat per mole of injected SDS at 25 °C is represented as a function of SDS concentration in the cell. Given the sigmoidal shape of the thermogram, the cmc° value can be identified as the inflection point. Inset shows the first derivative plot, which reaches a minimum at the inflection point allowing the accurate estimation of the cmc° value (B) The same procedure was repeated at different temperatures: 15 °C (purple), 22 °C (blue), 25 °C (light blue), 35 °C (green), 45 °C (light green), 55 °C (orange) and 65 °C (red)

It can be observed that the inflection point moves to higher concentration values when increasing the temperature (Fig. 3B).

Fig. 4 represents the obtained cmc° values as a function of temperature for both, C_12_E_10_ and SDS. It shows that cmc° values were always higher for SDS, and that the *cmc* temperature dependence for C_12_E_10_ is insignificant when compared with that for SDS.

**Fig. 4.**
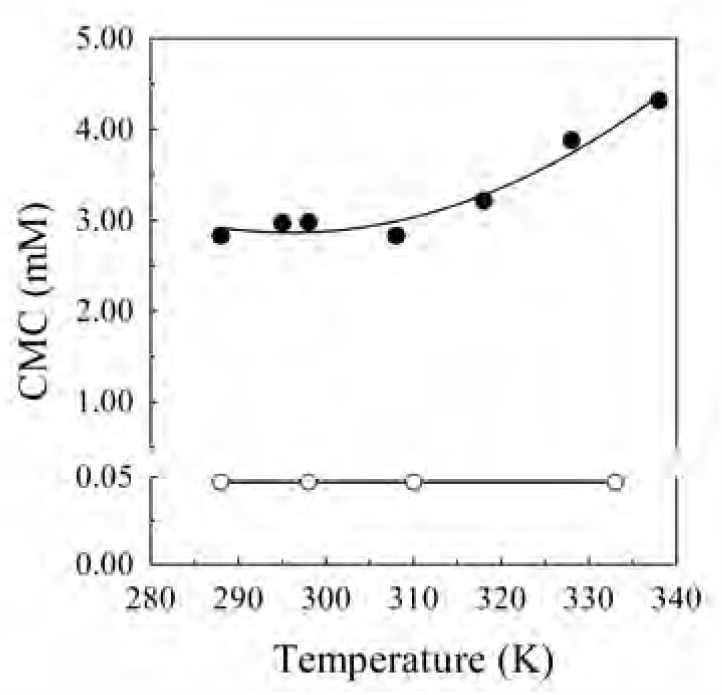
Dependence of the cmc° of SDS and C_12_E_10_ on temperature. The cmc° values obtained from Fig 2 and 3 for the titration of the buffer with each detergent were represented as a function of temperature. Continuous line represents empirical polynomial functions fitted to the experimental data. According to the Akaike criterion, the equations of best fit were: cmc°_C12E10_ = 47μM and cmc°_SDS_ = 0.0009 mM K^−2^ T^2^ - 0.5072 mM K^−1^ T + 77.317 mM, where T is the absolute temperature.

### 3.2. Determination of the composition of the micellar phase in the ATPase reconstitution medium

The composition of the micellar phase where the assayed CopAs were reconstituted is defined by the mole fractions of its components:

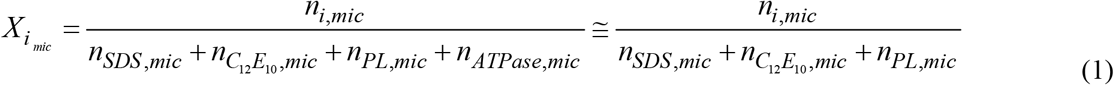

where *X*_i,mic_ is the mole fraction of the *i* component in the micelles (*i* may be phospholipid, detergent or protein); and *n* refers to the total amount of substance (number of moles).

Two assumptions were made for the approximation in the last member of Eq. (1): The first one is that the number of moles of protein in the micelle is much lower than that corresponding to amphiphiles (n_ATPase,mic_ << n_PL_). The other is that phospholipids are present exclusively in the micellar phase (*n*_*PL,mic*_*= n*_*PL,total*_), due to the very low critical micellar concentration values [24].

Considering that all the species (micelles and soluble amphiphiles) are in the same volume (which is assumed to be equal to the aqueous phase volume), and that amphiphiles distribute between the aqueous and the micellar phases, Eq (1) can be expressed as:

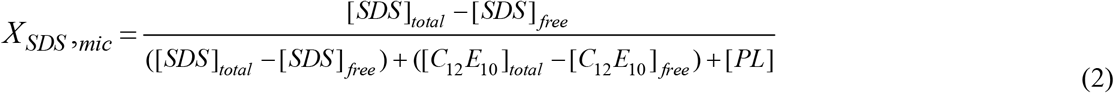

On the other hand, the free amphiphile concentration in the aqueous phase is related to the cmc of this amphiphile when it is the unique micelle component (cmc°) according to a Raoult-like relationship [25, 26].

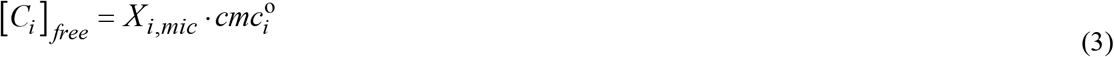

Finally, it holds that the sum of the mole fractions of all the amphiphiles in the micellar phase equals 1.

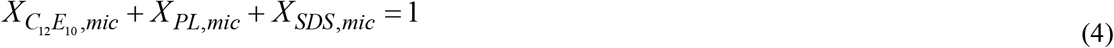

Despite the equations system (2), (3) and (4) having an analytical solution, it is very complex. Conversely, it can be easily solved numerically through an iterative procedure (Fig. 5).

**Fig. 5.**
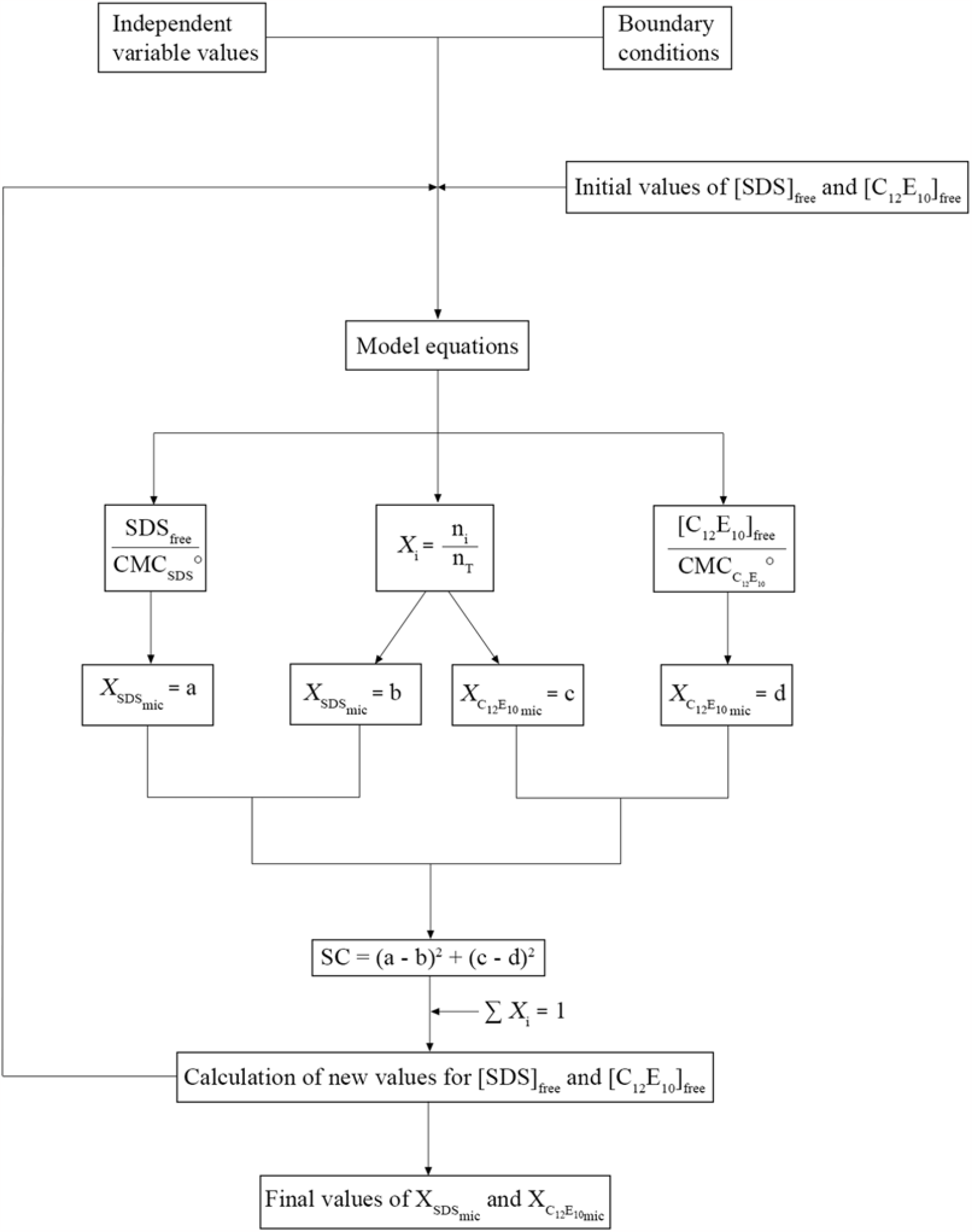
Iterative procedure for the determination of the micellar phase composition.

This procedure was applied to determine composition of the micellar phase when micelles of C_12_E_10_ and asolectin are supplemented with increasing quantities of SDS. Fig. 6 shows the relation between the proportion of each detergent in the amphiphile mixture as prepared (*X*_SDStotal_) and the same proportion inside the micelle once all the phases reach thermal and thermodynamic equilibria (*X*_SDS mic_). It can be observed that the enrichment of micelles is initially non favorable for SDS (because an important quantity of the SDS added remains in the aqueous phase). This behavior is more evident when the initial concentration of C_12_E_10_ is low. For example, to obtain a micellar phase that is equimolar in both detergents (*X*_SDS, mic_=0.5) we need to prepare a mixture with 81 % SDS (for [C_12_E_10_] = 0.36 mM) 73 % SDS (for [C_12_E_10_] = 0.72 mM), 65% SDS (for [C_12_E_10_] = 1.44 mM) or 52 % SDS (for [C_12_E_10_] = 20.5 mM).

**Fig. 6.**
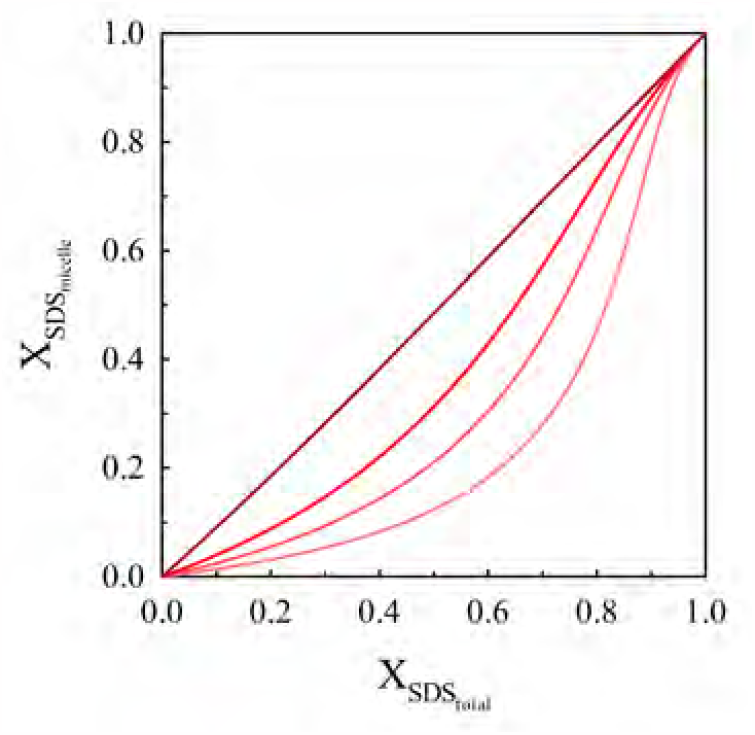
Relation between the mole fraction of SDS in the micelle and the fraction of the total amount of amphiphiles in the system. The mole fraction of SDS in the micelle was calculated as described in Fig 5 for the addition of increasing concentration of SDS to mixed micelles initially composed of C_12_E_10_ and asolectin with *X*_PL_= 0.15 and increasing concentrations of amphiphiles 0.43, 0.85, 1.7 and 24 mM (red lines with increasing color saturation). The fraction of SDS in the total amount of amphiphiles was calculated for each condition as a quotient between the number of moles of SDS added to the system and the number of moles of all the amphipiles (SDS, C_12_E_10_ and asolectin).

### 3.3. Inactivation of Cu(I) transport ATPases by sodium dodecyl sulfate

Purified *Lp*CopA was reconstituted in mixed micelles composed of 11 μM *Lp*CopA, 239 μM C_12_E_10_ in a medium containing 10 mM NaCl 25 mM Tris HCl pH 7 at 37°C. The enzyme was then diluted with 10 mM NaCl, 25 mM Tris HCl pH 7 at 37°C up to 0.8 μM CopA, with different asolectin/C_12_E_10_ concentration keeping constant the proportion between them, in the presence of increasing concentrations of SDS (Fig. 7A).

**Fig. 7:**
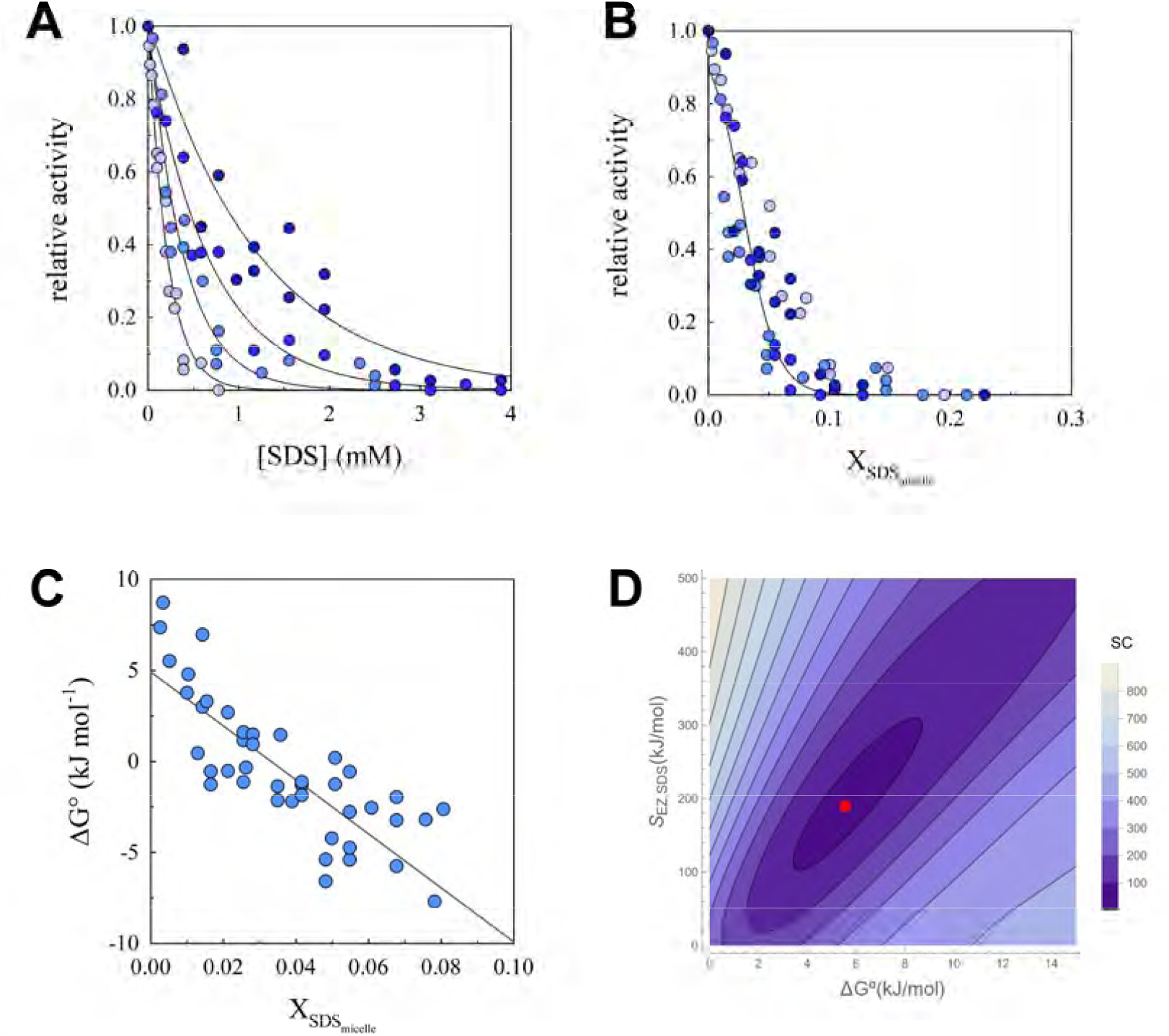
Effect of initial concentration of amphiphiles on the SDS-induced inactivation of *Lp*CopA. Purified *Lp*CopA (0.8 μM) was reconstituted in mixed micelles of asolectin and C_12_E_10_ (*X*_PL_= 0.15) with increasing concentrations of these amphiphiles 0.85, 12, 16 and 24 mM (blue dots with increasing color saturation). Samples were incubated with increasing concentrations of SDS for 5 min at 37°C, then the ATPase activity was measured as describe in materials and methods and represented as a function of the total concentration of SDS **(A)**, or of the mole fraction of SDS in the micelle **(B)**. Continuous lines are exponential fits to each individual set of experiments **(A)**, or the fit of Eq (9) to the full set **(B)**. Panel **(C)** shows the ΔG° values for the inactivation process calculated using Eq. (6) and (7) and represented as a function of *X*_SDS,mic_. **(D)** Contour plot showing the susceptibility coefficient as a function of ΔG° for different levels of the sum of squares corresponding to the fit of Eq. (9) to the full set of experimental data shown in panel **B**. The red dot indicates the best fit parameter values.

It can be observed that SDS produce the inactivation of the enzyme in all conditions, but it is required a higher SDS concentration to obtain the same degree of inactivation when the concentration of phospholipids and C_12_E_10_ is higher.

If we represent this data now as a function of the mole fraction of SDS in the micelle it can be observed that the three inactivation curves in Fig. 7A become a unique curve (Fig. 7B). This effect is not observed when representing the same data as a function of the total proportion of SDS in the amphiphile mixture (Fig S1A). It is worth nothing that in this case of the inactivation curves obtained with higher initial concentration of amphiphiles results are similar when represented as a function of *X*_SDS,mic_ or *X*_SDS, total_, but the curve corresponding to the lower amphiphile concentration only reaches the global trend when represented as a function of *X*_SDS, mic_ because in this condition there is an considerable difference between *X*_SDS, mic_ and *X*_SDS, total_ (Fig S1B). This is a very important result, because many structural studies on membrane proteins require reconstituting the protein with the lowest concentration of amphiphile compatible with function to avoid background interference [27].

Removing SDS from a full inactivated enzyme preparation (by dialysis against the enzyme incubation medium without SDS) results in the full recovery of the ATPase activity, indicating that the inactivation process is reversible.

To analyze this data we propose a simple model involving the transition between active (A) and inactive (I) enzyme molecules in equilibrium:

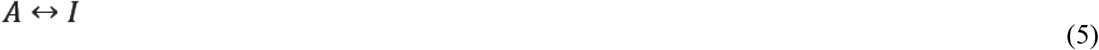

being the equilibrium constant

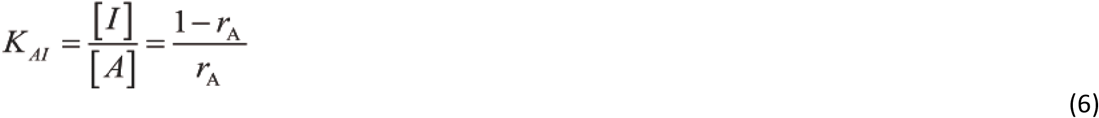

with *r*_A_ = relative ATPase activity. The Gibbs free energy for the transition between Active and Inactive enzyme, both at the reference state (hypothetical 1M ideal solutions) will be:

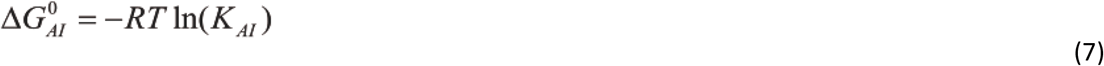

As can be observed in Fig. 7C, a linear dependence on *X*_SDS,mic_ can be observed. The slope of this relation represents the derivative of the Gibbs free energy for the inactivation process in the reference state, with respect to the mole fraction of SDS in the micellar phase. We can denote the positive value of this derivative as a measure of the susceptibility of the enzyme to SDS (*s*_Ez,SDS_), because it represents how much the enzyme inactivation equilibrium is shifted by an infinitesimal increase in the proportion of SDS in the micellar phase.

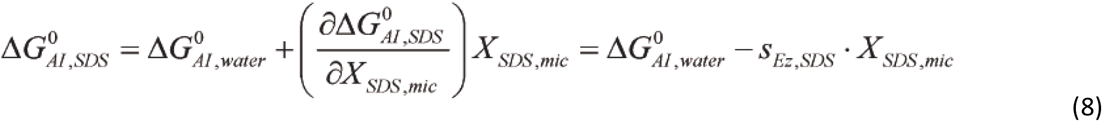

Combining Eq (6), (7) and (8), we obtain an expression of the relative activity (*r*_A_) as a function of the mole fraction of SDS in the micelle (*X*_SDS,mic_)

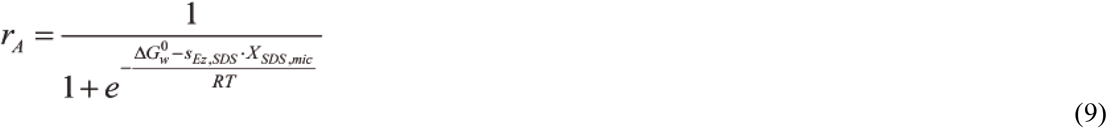

Continuous line in Fig 7B shows that the model represented by Eq (9) adequately describes the behavior of the experimental data. On the other hand, the contour plots for the sum of squares are indicative that there is a positive statistical correlation between the parameters of Eq (9).

It is easy to show that the mole fraction of SDS in the micelle producing half inactivation (*r*_*A*_*=0*.*5*) correspond to:

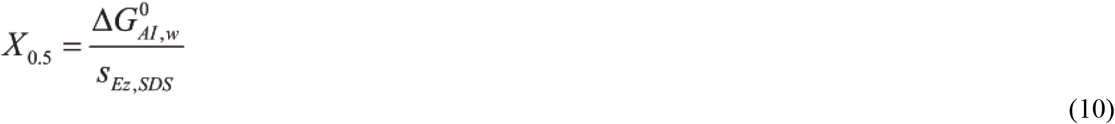

Fig. 8 shows the comparative effect of SDS inactivating two Cu(I) ATPases in the same buffer. *Lp*CopA (measuring the activity at 37°C) and *Af*CopA (measuring the activity at 70°C). Given the strong difference in the cmc value for SDS at 70°C it is evident that we will need to add much more SDS in the case of *Af*CopA to obtain the same micellar composition at 70°C that those obtained for *Lp*CopA at 37°C.

**Fig. 8.**
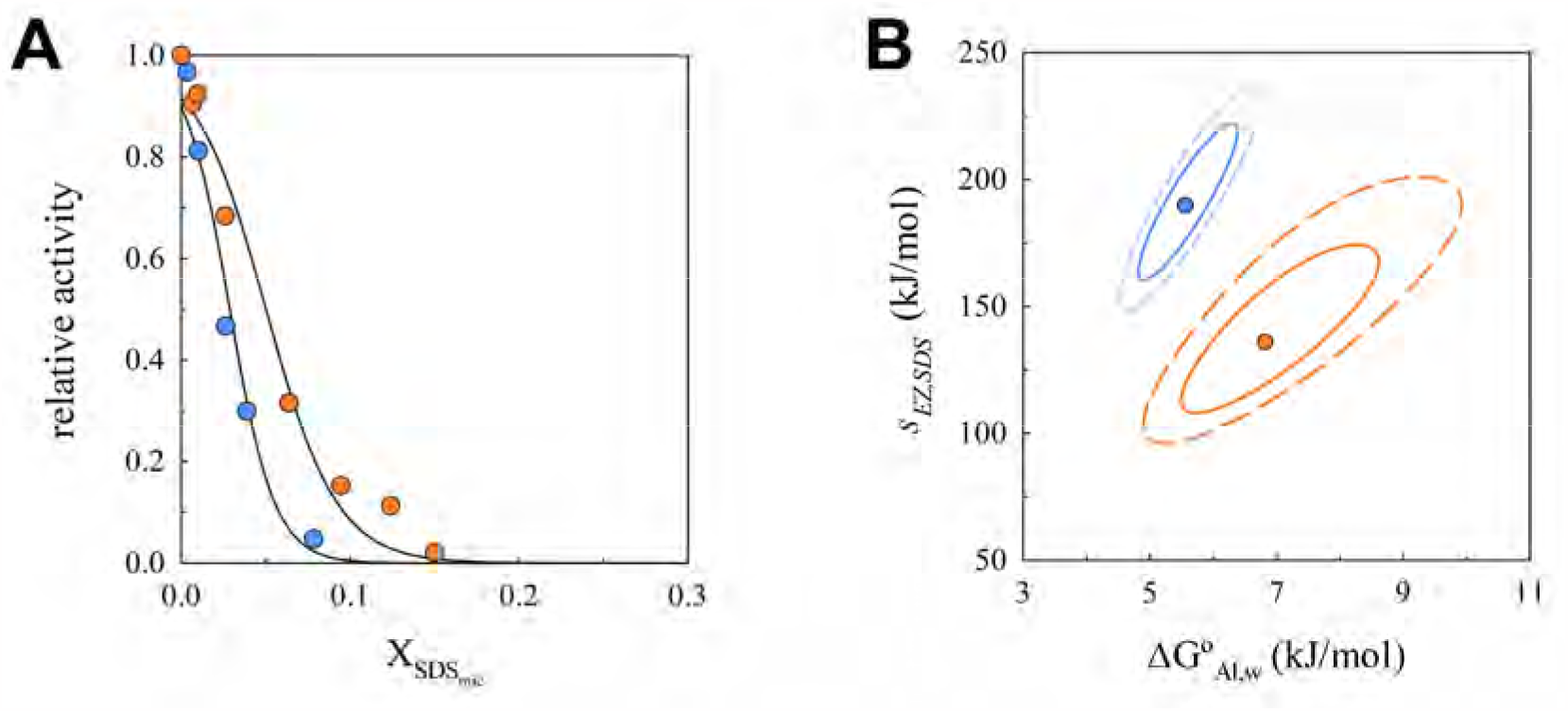
Inactivation by SDS of two Cu(I) transport ATPases with different thermal adaptation. Purified *Af*CopA (orange circles) and *Lp*CopA (blue circles) were reconstituted in mixed micelles of asolectin and C_12_E_10_. Samples were incubated with increasing concentrations of SDS for 5 min at 37°C (*Lp*CopA) and 70°C (*Af*CopA), then the ATPase activity was measured as describe in materials and methods and represented as a function of the mole fraction of SDS in the micelle **(A)**. Continuous lines are the graphical representation of Eq (9) fitted to the experimental data. Panel **B** shows the 68% confidence region (continuous lines), and 95 % confidence region (dashed lines). The best fit parameter values are shown in Table I.

**Table I.** Model parameters for the inactivation of Cu(I) ATPases by SDS

| Enzyme | Temperature (K) | $\Delta G^{\circ}_{Al,w}$ (kJ/mol) | $s_{Ez,SDS}$ (kJ/mol) | $X_{0.5}$ |
| --- | --- | --- | --- | --- |
| *AfCopA* | 343 | $6.8 \pm 0.9$ | $135 \pm 18$ | 0.05 |
| *LpCopA* | 310 | $5.5 \pm 0.6$ | $190 \pm 20$ | 0.03 |

As a control we explore the circular dichroism and tryptophan fluorescence spectra, two of the main structural parameters used to characterize the native state of the protein, at a SDS mole fraction producing full inactivation of the enzyme. It was observed that the circular dichroism spectrum remains unchanged, and only a 3% decrease in the fluorescence intensity was observed.

It can be observed that the inactivation free energy changes are similar for both proteins (each one at its working temperature) and correspond to an equilibrium constant for inactivation of about 0.11 indicating that the equilibrium in the aqueous/micellar medium is highly shifted towards the active form. Conversely, the value of the susceptibility to SDS is significantly higher for *Lp*CopA indicating that *Af*CopA is more resistant to SDS, agreeing with its thermophilic nature. Further studies are needed to verify if this behavior can be extended to other enzymes.

## 4. Concluding remarks

In this work we have explored the effect of SDS at sub-denaturing concentrations on the ATPase activity of a hyper-thermophilic (*Af*CopA) and a mesophilic (*Lp*CopA) Cu(I) transport ATPases reconstituted in mixed micelles of phospholipids and a non-denaturing detergent. An iterative procedure was developed to evaluate the partition of SDS between the aqueous and the micellar phases. This procedure made it possible to determine that a very different composition of micelles is obtained when the mixture is prepared with the same proportions of amphiphiles but a different amount of non-denaturing amphiphiles.

Inactivation of *Lp*CopA by incubation with SDS in the presence of different amounts of phospholipids led to the conclusion that the mole fraction of SDS in the micelle is the compositional variable that should be used to analyze these data. A simple model involving active and inactive enzyme molecules in equilibrium was proposed to give process (in the reference states) and its derivative respect to the mole fraction of SDS in the micellar phase. This last parameter gives a measure of the susceptibility of the enzyme to SDS.

Comparing the inactivation behavior of *Af*CopA and *Lp*CopA, we found that the inactivation free energy changes were similar for both proteins, but the value of the susceptibility to SDS was significantly lower for the hyperthermophilic ATPase.

This result suggests an inverse relation between thermophilicity and susceptibility to SDS.

## Acknowledgements

The authors are grateful to Dr J. Jeremias Incicco for critical reading of the manuscript. This work was financed with grants from Universidad de Buenos Aires (UBACyT 306BA), CONICET (PIP 3266CO) and agencia I+D+I (PICT 2019 - 02768).

## Abbreviations

*Af*CopA: Cu(I) transport ATPase from *Archaeoglobus fulgidus*
C_12_E_10_: polyoxyethylene 10 lauryl ether
*cmc*: critical micelle concentration
*Lp*CopA: Cu(I) transport ATPase from *Legionella pneumophila*
PL: phospholipids
SC: sum of squares
SDS: sodium dodecyl sulfate
*s*_Ez,SDS_: susceptibility of a given enzyme to SDS
*X*: mole fraction

## References

[1] A.A. Recoulat Angelini, M.A. Placenti, N.A. Melian, M.L. Sabeckis, N.I. Burgardt, R.M. González Lebrero, E.A. Roman, F.L. González Flecha, Cu(I)-Transport ATPases Molecular Architecture, Catalysis and Adaptation to Extreme Environments, Adv Med Biol, 180 (2021) 65–130

[2] N. Salustros, C. Grønberg, N.S. Abeyrathna, P. Lyu, F. Orädd, K. Wang, M. Andersson, G. Meloni, P. Gourdon, Structural basis of ion uptake in copper-transporting P1B-type ATPases, Nat Commun, 13 (2022) 5121.DOI: 10.1038/s41467-022-32751-w

[3] F. Cymer, G. von Heijne, S.H. White, Mechanisms of Integral Membrane Protein Insertion and Folding, J Mol Biol, 427 (2015) 999–1022.DOI: https://doi.org/10.1016/j.jmb.2014.09.014

[4] M.F. Pignataro, M.M. Dodes-Traian, F.L. González-Flecha, M. Sica, I.C. Mangialavori, J.P.F.C. Rossi, Modulation of Plasma Membrane Ca2+-ATPase by Neutral Phospholipids: Effect of the micelle-vesicle transition and the bilayer thickness J Biol Chem, 290 (2015) 6179–6190.DOI: 10.1074/jbc.M114.585828

[5] J.A. Reynolds, C. Tanford, Binding of dodecyl sulfate to proteins at high binding ratios. Possible implications for the state of proteins in biological membranes, Proc Natl Acad Sci U S A, 66 (1970) 1002–1007.DOI: 10.1073/pnas.66.3.1002

[6] E.A. Roman, F.L. González Flecha, Kinetics and thermodynamics of membrane protein folding, Biomolecules, 4 (2014) 354–373.DOI: 10.3390/biom4010354

[7] D.E. Otzen, J.N. Pedersen, H.Ø.Rasmussen, J.S. Pedersen, How do surfactants unfold and refold proteins?, Adv Coll Interf Sci, 308 (2022) 102754.DOI: https://doi.org/10.1016/j.cis.2022.102754

[8] D.E. Otzen, P. Sehgal, P. Westh, α-Lactalbumin is unfolded by all classes of surfactants but by different mechanisms, J Coll Interf Sci, 329 (2009) 273–283.DOI: https://doi.org/10.1016/j.jcis.2008.10.021

[9] S. Baird, S.M. Kelly, N.C. Price, E. Jaenicke, C. Meesters, D. Nillius, H. Decker, J. Nairn, Hemocyanin conformational changes associated with SDS-induced phenol oxidase activation, Biochim Biophys Acta, 1774 (2007) 1380–1394.DOI: https://doi.org/10.1016/j.bbapap.2007.08.019

[10] H. Mu, S.-M. Zhou, Y. Xia, H. Zou, F. Meng, Y.-B. Yan, Inactivation and Unfolding of the Hyperthermophilic Inorganic Pyrophosphatase from Thermus thermophilus by Sodium Dodecyl Sulfate, in: Int. J. Mol. Sci., vol. 10, 2009, pp. 2849–2859.

[11] K. Xia, S. Zhang, B. Bathrick, S. Liu, Y. Garcia, W. Colón, Quantifying the Kinetic Stability of Hyperstable Proteins via Time-Dependent SDS Trapping, Biochemistry, 51 (2012) 100–107.DOI: 10.1021/bi201362z

[12] A.K. Mandal, W.D. Cheung, J.M. Argüello, Characterization of a thermophilic P-type Ag^+/Cu+^-ATPase from the extremophile Archaeoglobus fulgidus, J Biol Chem, 277 (2002) 7201–7208.DOI: 10.1074/jbc.M109964200

[13] M.A. Placenti, E.A. Roman, F.L. González Flecha, R.M. González-Lebrero, Functional characterization of Legionella pneumophila Cu+ transport ATPase. The activation by Cu+ and ATP, Biochim Biophys Acta - Biomembr, 1864 (2022) 183822.DOI: https://doi.org/10.1016/j.bbamem.2021.183822

[14] E.A. Roman, J.M. Argüello, F.L. González Flecha, Reversible unfolding of a thermophilic membrane protein in phospholipid/detergent mixed micelles, J. Mol. Biol., 397 (2010) 550–559.DOI: 10.1016/j.jmb.2010.01.045

[15] C.N. Pace, F. Vajdos, L. Fee, G. Grimsley, T. Gray, How to measure and predict the molar absorption coefficient of a protein, Protein Sci, 4 (1995) 2411–2423.DOI: 10.1002/pro.5560041120

[16] S.A. Martínez Gache, A.A. Recoulat Angelini, M.L. Sabeckis, F.L. González Flecha, Improving the stability of the malachite green method for the determination of phosphate using Pluronic F68, Anal Biochem, 597 (2020) 113681.DOI: https://doi.org/10.1016/j.ab.2020.113681

[17] A.A. Recoulat Angelini, S.A. Martínez Gache, M.L. Sabeckis, N.A. Melian, F.L. González Flecha, On the role of citrate in 12-molybdophosphoric-acid methods for quantification of phosphate in the presence of ATP, New J. Chem., 46 (2022) 12401–12409.DOI: 10.1039/d2nj00943a

[18] S.-C. Tso, F. Mahler, J. Höring, S. Keller, C.A. Brautigam, Fast and Robust Quantification of Detergent Micellization Thermodynamics from Isothermal Titration Calorimetry, Anal. Chem., 92 (2020) 1154–1161.DOI: 10.1021/acs.analchem.9b04281

[19] G. Kemmer, S. Keller, Nonlinear least-squares data fitting in Excel spreadsheets, Nat Protoc, 5 (2010) 267–281.DOI: 10.1038/nprot.2009.182

[20] H. Akaike, A Bayesian analysis of the minimum AIC procedure, Ann. I. Stat. Math., 30 (1978) 9–14.DOI: 10.1007/BF02480194

[21] G.E.P. Box, J.S. Hunter, G.W. Hunter, Statistics for Experimenters: Design, Innovation, and Discovery, 2nd ed., Wiley-Interscience, Hoboken, NJ,, 2005.

[22] V. Levi, A.M. Villamil Giraldo, P.R. Castello, J.P.F.C. Rossi, F.L. González Flecha, Effects of phosphatidylethanolamine glycation on lipid–protein interactions and membrane protein thermal stability, Biochem J, 416 (2008) 145–152.DOI: 10.1042/bj20080618

[23] D.N. LeBard, B.G. Levine, R. DeVane, W. Shinoda, M.L. Klein, Premicelles and monomer exchange in aqueous surfactant solutions above and below the critical micelle concentration, Chem Phys Letters, 522 (2012) 38–42.DOI: https://doi.org/10.1016/j.cplett.2011.11.075

[24] F. Scollo, C. Tempra, F. Lolicato, M.F.M. Sciacca, A. Raudino, D. Milardi, C. La Rosa, Phospholipids Critical Micellar Concentrations Trigger Different Mechanisms of Intrinsically Disordered Proteins Interaction with Model Membranes, J Phys Chem Letters, 9 (2018) 5125–5129.DOI: 10.1021/acs.jpclett.8b02241

[25] C. Tanford, Theory of micelle formation in aqueous solutions, J. Phys. Chem., 78 (1974) 2469–2479.DOI: 10.1021/j100617a012

[26] C. Tanford, The hydrophobic effect: Formation of micelles and biological membranes, John Wiley & Sons Inc, 1980.

[27] A.J. Miles, B.A. Wallace, Circular dichroism spectroscopy of membrane proteins, Chem. Soc. Rev., 45 (2016) 4859–4872.DOI: 10.1039/c5cs00084j

